# Nucleus accumbens neuron subtype translatome signatures in socially stressed females

**DOI:** 10.1101/2025.06.01.656319

**Authors:** Gautam Kumar, Daniela Franco, Mahashweta Basu, Jimmy Olusakin, Rianne Campbell, Seth A. Ament, Megan E. Fox, Mary Kay Lobo

## Abstract

Major depressive disorder (MDD) disproportionately affects women, yet the molecular mechanisms underlying female vulnerability remain poorly understood. The nucleus accumbens (NAc), a reward circuitry hub, is central to stress- and depression-related pathology. While NAc medium spiny neuron (MSN) transcriptional adaptations have been characterized in stressed male rodents, much less is known about female adaptations. To address this, we exposed female D1-Cre-RiboTag and A2A-Cre-RiboTag mice to chronic witness defeat stress (CWDS) and classified them as high- or low-social interactors. Ribosome-associated mRNA was isolated from MSN subtypes and analyzed by RNA sequencing, differential gene expression analysis (DEG), and weighted gene co-expression network analysis (WGCNA). Cross-species consensus WGCNA with male mouse and human transcriptomic datasets was performed to identify conserved, sex-specific signatures. We detected 9 differentially expressed genes (DEGs) in D1-MSNs and 630 in A2A-MSNs (FDR < 0.05). In D1-MSNs, DEGs were primarily upregulated in low interactors and enriched for energy homeostasis and cell adhesion. A2A-MSN DEGs included upregulated structural genes and downregulated neurotransmission-related genes. WGCNA identified 9 significant D1- and 5 significant A2A-MSN modules, with the most impacted enriched for PI3K-Akt-mTOR signaling and regulated by Nf1. Consensus analysis revealed a stress-associated, sex- and subtype-specific module enriched for PI3K-Akt-mTOR signaling conserved across mice and humans. These findings reveal female-specific MSN transcriptional adaptations to chronic social stress and implicate PI3K-Akt-mTOR signaling as a convergent molecular pathway. By highlighting MSN subtype-specific vulnerabilities, this work suggests potential therapeutic targets for alleviating stress-induced pathology in women with MDD.

## Introduction

Major depressive disorder, (MDD), is expected to become the foremost cause of disease burden worldwide by 2030 [1]. Treatment has limited success, with 30% of patients resistant or relapsing [2]. To uncover the mechanisms underlying vulnerability to MDD, researchers use chronic stress paradigms to model disrupted affective behaviors. Chronic social stress paradigms have face validity and better mirror the psychosocial nature of human stress [3–9]. These paradigms, however, are often limited to male-only use[10]. Given sex differences in MDD symptomatology, antidepressant response, and increased incidence in women, female subjects are crucial for understanding MDD neuropathology [10]. Social stress models employing female subjects have emerged in recent years [8, 12, 14]. One social stress paradigm, chronic witness defeat stress (CWDS) adapts chronic social defeat stress (CSDS) by exposing mice to witnessed - rather than physical - defeat allowing modeling of affective dysfunction in females [8, 11]. Unlike other alternative models, employing female aggressors, male odorants or chemogenic manipulation, CWDS models the psychological aspects of social trauma, making it more translatable, as sources of human stress tend to be interpersonal/emotional [12–14].

Understanding mechanisms in brain regions mediating depression symptoms is essential for novel therapies. One such region is the nucleus accumbens (NAc), a key integration center in the motivational circuitry [7, 15, 16]. MDD patients show reduced NAc volume and blunted activation, while deep brain stimulation of the NAc alleviates symptoms in resistant cases, driving interest in NAc-targeted treatments [17]. Medium spiny neurons (MSNs) make up over 90% of NAc neurons and include D1- and D2-type subtypes distinguished by dopamine receptor expression and other GPCRs like adenosine 2a receptor (A2A) [18, 19]. Activation of D1-MSNs promotes reward behavior, while inhibition increases vulnerability to anhedonia [20, 21]. Increased D1-MSN intrinsic excitability with reduced synaptic input, and enhanced D2/A2A-MSN synaptic input, associate with social avoidance and stress susceptibility [20]. However, how sex differences in these subtypes contribute to MDD remains unclear.

Using female CWDS and celltype-specific RNA-seq via D1-Cre-RiboTag and A2A-Cre-RiboTag mice, we profiled NAc MSN subtype-specific translatomes. We identified differentially expressed genes and co-expression networks in D1- and A2A-MSNs of low and high SI females. We then used publicly available transcriptomics data to identify consensus networks in male and female mice and human MDD patients, identifying unique sex-specific molecular signatures relevant to MDD [22, 23]. Given the chronic, relapsing course of MDD, this study may reveal molecular targets for MDD and other stress-related psychiatric disorders.

## Materials and Methods

### Animals

Experiments followed Institutional Animal Care and Use Committee guidelines at University of Maryland School of Medicine (UMSOM). Mice were given ad libitum food/water and housed in a 12:12 h light:dark cycle in the UMSOM vivarium. Homozygous Rpl22HA (“RiboTag”, RT) mice crossed with D1-Cre (line FK150)/A2A-Cre (line KG 139) mice generated female D1-Cre-RT & A2A-Cre-RT mice used for subtype-specific RNA-seq [24]. All transgenic mice were on a C57Bl/6 background and 8-10 weeks old at the start. Male CD-1 retired breeders (>4 months, Charles River) served as CWDS aggressors; male C57BL6 conspecifics were intruders.

### Chronic Witness Defeat Stress (CWDS)

CWDS was performed as described [14]. Female experimental mice, housed behind a plexiglass divider in a cage, witnessed a male CD-1 defeat a male intruder for 10 min. This occurred between 9:00-14:00 ZT each day. After each witness episode the witness mouse was then housed opposite a novel CD-1 for 24-hour sensory interaction. This repeated for 10 days with a novel CD-1 resident each day. Control mice were pair-housed with sex-matched conspecifics, separated by a divider, for 10 days. After the last witness episode mice were singly housed through the three-chamber social interaction on day 11 followed by tissue collection on day 12.

### Three-chamber Social Interaction Test

24 hours post-defeat, mice underwent 3 chamber social interaction (3ChSI) testing between 9:00-14:00 ZT. Mice started in the center of a three-chambered arena (60 × 40 cm), with empty cups in peripheral chambers and explored for 5 minutes. A sex-matched conspecific was then placed under one cup, and the same mouse explored for 5 more minutes. Time spent in the social chamber was video-tracked (TopScan Lite, CleverSys) to assess stress phenotype. Classification was based on the average social chamber time of controls (both D1- and A2A-Cre-RT controls; ∼144s). Mice spending <144s were classified as low social interactors while ≥144s were classified as high social interactors. One-way ANOVA with Tukey’s post hoc test assessed stress group effects.

### Tissue collection & RNA isolation

24h after social interaction testing at 9:00-14:00 ZT, four 14 gauge punches containing the entire NAc that includes both core and shell were collected from D1-Cre-RT and A2A-Cre-RT mice and snap frozen at −80C. Polyribosomes were immunoprecipitated as previously described [25], each sample pooled from four mice with similar SI ratios. Homogenates were incubated with 5 μl anti-HA antibody (Biolegend 901515) at 4°C overnight with rotation. Samples were then incubated with 400 uL of protein G magnetic beads (Life Technologies, Carlsbad, CA, USA #100.09D) overnight at 4°C with rotation. Beads were washed in a magnetic rack with high-salt buffer. RNA was extracted with a DNase step (Qiagen, Germantown, MD, USA) using the RNeasy mini kit (qiagen) by adding RLT and following manufacturer instructions. RNA concentration and quality were assessed via Bioanalyzer (Agilent).

### Library preparation and analysis

Samples with RNA integrity number >8 were used and pooled into biological quintuplicates for sequencing at the Maryland Genomics core of the UMSOM Institute for Genome Sciences (IGS) and processed as described previously [26]. Libraries were prepared from 10 ng of RNA from each sample using the SMART-Seq v4 kit (Takara). Samples were sequenced using NovaSeq6000 with 100 bp paired-end reads. Reads were aligned to the mouse genome (Mus musculus, GRCm38) using HISAT2 and counted in coding regions using HTSeq (GEO: GSE315159).

We used two approaches to characterize transcriptomic alterations. First, edgeR’s likelihood ratio test identified DEGs by stress group in MSN subtypes. Genes with False Discovery Rate (FDR) < 0.05 were considered differentially expressed. Gene Ontology (GO) enrichment was performed using R packages ‘gprofiler2’ and ‘clusterProfiler’ and the online tool ‘DAVID’. We next used weighted gene co-expression network analysis (WGCNA) to identify networks of co-expressed genes in D1- and D2-MSNs. ‘blockwiseModules()’ identified signed modules using biweight midcorrelation, power = 12, and minimum module size = 25. Module membership scores were calculated by correlating gene expression with module eigengene. Hub genes were defined as those with highest membership scores. Using the R package ‘limma’, differentially regulated modules were identified using module eigengenes in a linear model comparing stress groups. Significant modules were characterized using the same enrichment tools.

WGCNA modules were classified into four categories based on hub gene properties and enrichment terms: structural, synaptic, protein synthesis and mitochondrial. Modules dominated by genes and terms linked to morphogenesis, junction/projection formation, cytoskeletal dynamics and ECM interactions were classified as structural. Modules with genes and terms heavily involved in synaptic response, receptor trafficking and signal transduction were classified as synaptic. Protein synthesis modules were composed of genes encoding ribosomal and proteasomal subunits, transcription factors and elongation factors and genes involved in protein processing. Finally, mitochondrial modules were largely composed of respiratory complex components. In cases where properties of modules overlapped between multiple categories, modules were classified into the group that most closely aligned with the properties of modular hub genes.

RNA-seq data from male D1- and D2-Cre-RiboTag mice and human male and female NAc postmortem tissue were obtained from GEO [23, 27]. Male mouse reads were downloaded and processed using RSubread (GRCm39); human read counts were downloaded processed.

### Consensus module detection and analysis

Human genes were mapped to mouse orthologs using the R package ‘orthogene’. Genes with low counts were filtered using edgeR’s *filterByExpr*, and normalization used variance stabilizing transformation. Consensus modules were constructed using all datasets (power = 8 and minimum module size = 30). ‘Limma’ identified differentially regulated modules in each dataset.

## Results

### Social stress in females induces differentially enriched translatomes in nucleus accumbens neuron subtypes

Female D1-Cre-RT and A2A-Cre-RT mice underwent 10 days of CWDS and three chamber social interaction test on day 11 (Fig 1A). High- and low-social interactors (SI) were classified based on control mice’s average time in the social chamber (∼144 s). This approach captures the natural range of social behavior exhibited by animals and enables comparison across groups. ANOVA confirmed significant differences in interactions times between stress groups (D1: p-value < 0.05, F(2,57) = 45.09; A2A: p-value < 0.05, F(2,58) = 25.12, Fig 1B). On day 12 NAc tissue was isolated, HA-tagged polyribosomes immunoprecipitated from D1- or A2A-MSNs, and ribosome-associated RNA isolated and sequenced.

**Fig 1.**
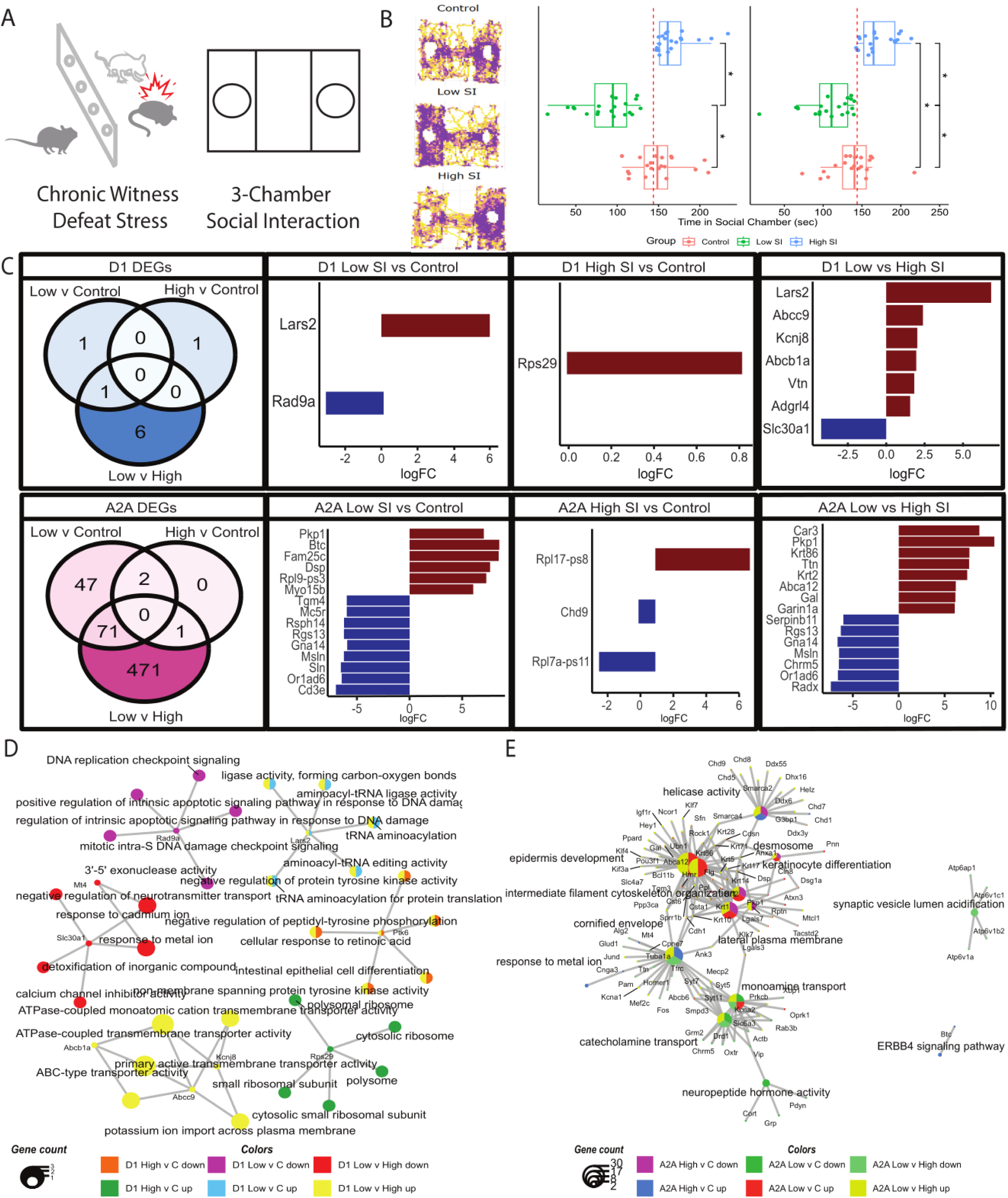
A) Schematic of experimental design. D1-Cre-RT and A2A-Cre-RT female mice were made to undergo the witness defeat paradigm for 10 days (10 minutes/day). 24 hours following the last defeat, the mice went through the 3 Chamber Social Interaction Test (3ChSI) after which they were sacrificed 24h later. The average time control mice spent in the social chamber (D1- and A2A-Cre-RT controls combined) was used to classify mice has high social interactors (higher than control average) or low social interactors (lower than control average). B) (left) Heatmap displaying localization of low social interactors (SI), high SI and control mice during the 3ChSI test. Areas highlighted in purple correspond with locations mice spend more time in while areas in yellow correspond with areas mice spent less time in. (right) Boxplots showing the time mice from each group spent in the interaction chamber (for each MSN subtype; one-way ANOVA: D1: p-value < 0.05, F(2,57) = 45.09; A2A: p-value < 0.05, F(2,58) = 25.12). The red dotted line represents the control group average, which was used to classify mice as low or high SI. C) (left to right) Venn diagrams visualizing the overlapping DEGs across stress groups in (top) D1- and (bottom) A2A-Cre-RT mice, bar plots of top differentially expressed genes (DEGs) between the stress groups in (top) D1- and (bottom) A2A-Cre-RT mice. D) Enrichment results for D1-MSN DEGs with the genes and terms color coded based on the DEG list they represent. E) Enrichment results for A2A-MSN DEGs with the genes and terms color coded based on the DEG list they represent.

We found 9 differentially expressed genes (DEGs) in D1-MSN and 630 in A2A-MSN across stress groups (FDR < 0.05, Fig 1C, Fig S1A-B, Table S1). Seven D1-MSN DEGs differ between high and low SI, with one downregulated in low SI (Fig 1C, top & far right). The top three upregulated genes – *Lars2, Abcc9* and *Kcnj8* -participate in cellular responses to energy deficiency, including an ATP-sensitive inward-rectifying potassium (KATP) channel.

Other upregulated genes are involved in adhesion/migration. Enrichment analysis of upregulated low vs high SI female D1-MSN DEGs include terms such as “ATPase-coupled transmembrane transporter activity” and “potassium import across membrane” while downregulated DEGs are involved in ion transport (Fig 1D). Enrichment analysis also indicates increased expression of genes involved in translation in high SI females and DNA damage response in low SI mice (Table S2).

Due to the low D1-MSN DEG count, enrichment analysis was repeated using a nominal p-value 0.05 cut-off. Low SI upregulated D1-MSN genes are enriched for chromatin- and synapse-related terms; downregulated genes are enriched for apoptosis. High SI D1-MSN upregulated genes are enriched for translation and mitochondrial/respiration terms; downregulated genes involve structural development (Table S1, S3).

A2A-MSNs had more DEGs. In low-SI vs. control/high-SI females, upregulated genes were linked to junction formation, while downregulated genes were involved in synaptic transmission, particularly GPCR signaling (Fig 1C, bottom). In high SI vs control, two ribosomal subunit pseudogenes were differentially regulated. Enrichment analysis showed upregulated genes in low-SI mice were associated with structural development and transcription regulation (Fig 1E, Table S2). Downregulated genes in low SI vs control/high SI genes were enriched for synaptic processes, notably GPCR signaling (Fig 1E). Enrichment analysis using a nominal p-value cutoff produced similar results for all comparisons (Table S3). There is little correlation between the MSN subtypes in gene expression change (Fig S1C).

### Female social stress results in altered gene networks in nucleus accumbens neuron subtypes

Weighted Gene Co-expression Network Analysis (WGCNA) of female D1- and A2A-Cre-RT transcriptomes identified 43 D1-MSN and 21 A2A-MSN modules (Fig. 2A-B, 3A-B, S2A-B, Table S4), containing between 33–3263 and 131–3064 genes, respectively (Fig. 2C, 3C). 9 D1-MSN and 5 A2A-MSN modules were significantly associated with high- or low-social interaction (Fig. 2B-C, 3B-C, Table S4). Given that the samples were pooled, we constructed boxplots using the eigengenes of the significant modules across pooled samples to examine potential sources of variability. Our results for both the D1- and A2A-Cre-RT modules indicate that module score medians all center around 0 with similar interquartile ranges. Eigengene variability doesn’t indicate any kind of instability across pools and there is no indication of any individual pooled sample driving eigengene signal (Fig S3, S4).

**Fig 2.**
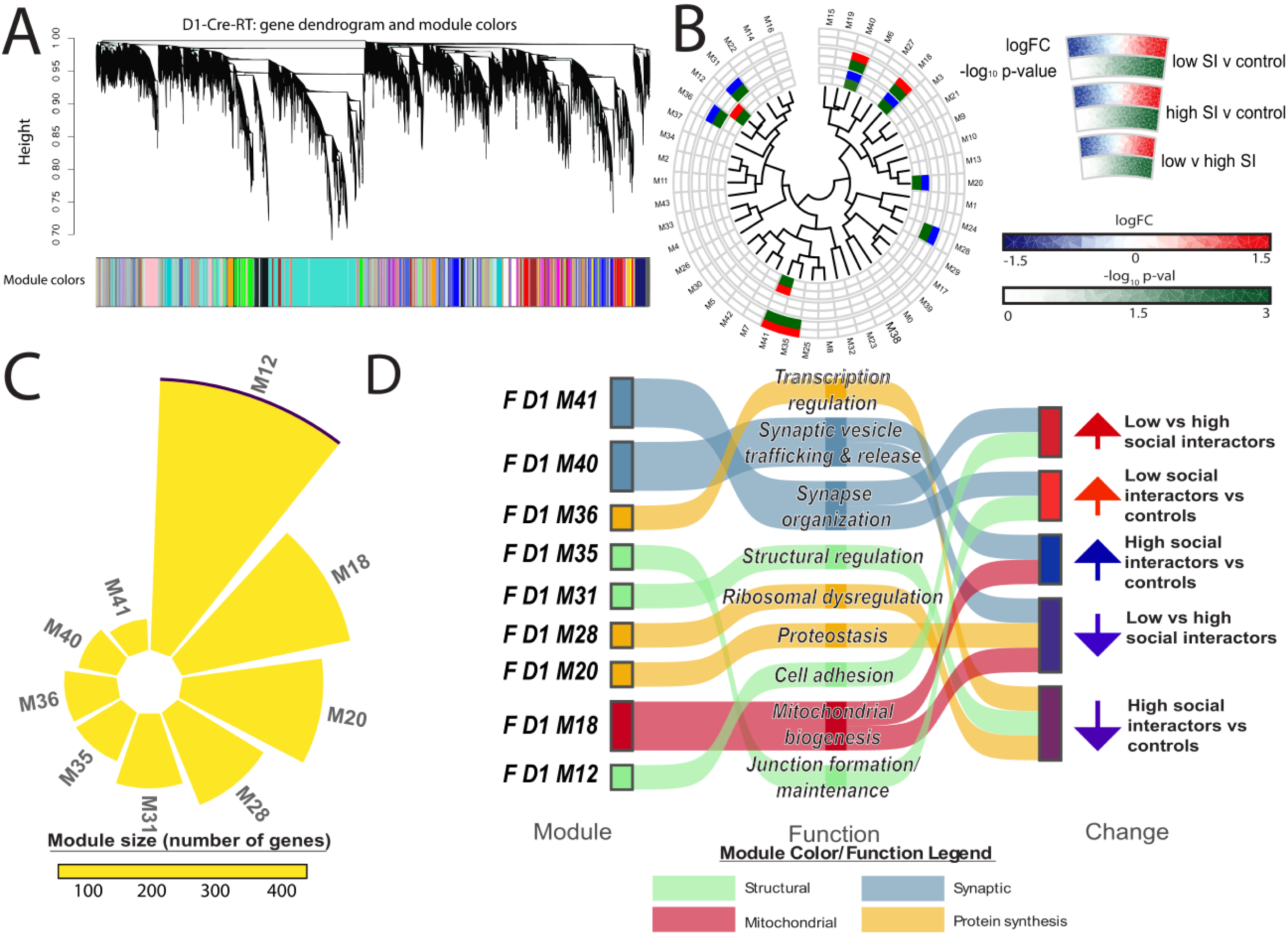
A) Dendrogram of genes from the female D1-Cre-RT data with WGCNA module assignment represented in the colored bar below. B) Circos plots of the significant D1-MSN modules with each ‘slice’ corresponding with one module. The plot is organized based on the module eigengene dendrogram, shown in the center, which positions related modules close to one another. Modules that aren’t significant are uncolored. C) Circular bar plots of D1-MSN modules significantly associated with high or low social interaction in female socially stressed mice. The proportion of the module that includes differentially expressed genes from the D1 high SI vs low SI comparison is highlighted in purple. D) Sankey plots of significantly differentially regulated modules in the D1-MSN of female socially stressed mice; modules are renamed based on their original WGCNA module number and functional categorization based on hub gene and enrichment analysis characterization.

The significant modules were characterized as described in the methods and placed into four categories: structural processes, protein synthesis, synapse/signaling and mitochondria related (Fig 2D, 3D, Table S5). WGCNA was also applied to chronically stressed male MSN subtypes and human NAc MDD datasets, identifying seven D1- and one A2A-MSN module associated with social interaction, and 12 human NAc modules significant by sex, condition, or both (Fig S3-S5, Table S6-9) [27].

Significant modules across female stress groups highlight the diverse effects of chronic social stress on MSN transcriptomes. While both female SI groups show differential regulation in D1-MSN modules, A2A-MSN modules are primarily affected in low SI females (Fig 2B, 3B). In contrast, male CSDS data show D1-MSN modules primarily impacted in low SI mice with one A2A-MSN module upregulated in both stress groups (Fig S5B, Fig S7B) [27]. Module preservation analysis demonstrated a lack of preservation between male and female mice, pointing to sex-specific transcriptional network architecture in these neurons and indicating that chronic stress impact distinct molecular mechanisms by sex (Fig S6, S8). This could also be indicative to the different types of stress-physical vs. emotional- and future studies are warranted to better understand these differences.”

Protein turnover modules, F D1 M20 and F A2A M3, are downregulated in low vs high SI (Fig 2D, 3D). F D1 M20 hub genes regulate transcription elongation, RNA degradation, and cytoskeletal dynamics, whereas F A2A M3 hub genes involve tRNA loading, RNA splicing, and cell adhesion (Fig 2D, 3D, Table S4). Enrichment results support these findings, with F D1 M20 genes enriched for elongation, ribosome, and proteasome terms, and F A2A M3 genes for localization and transport terms (Table S5). Multiple significant female modules highlight modulatory processes in protein synthesis and transport. The F D1 M28 module, downregulated in high SI vs controls, includes ribosomal subunit pseudogenes and stress response genes, (such as Rgs2, Manf, Isg15), suggesting ribosomal dysfunction (Fig 2D, Table S4-5). F D1 M36 is enriched for transcription regulation, Golgi function and RNA polymerase II activity while the F A2A M6 module is enriched for RNA polymerase II activity and ER-related terms (Fig 2D, 3D, Table S5). F D1 M36 is downregulated in high SI vs control, whereas F A2A M6 is upregulated in low vs high SI mice (Fig 2D, 3D). These results indicate reduced regulatory protein processes in D1-MSN of high SI females, but increased transcriptional regulation in A2A-MSNs of low SI females.

Structural regulation modules are exclusive to D1-MSNs. The F D1 M12 module, upregulated in low SI, includes hub genes involved in adhesion/migration, GPCR signaling, and stress response, with enrichment terms involving adhesion/migration and cytokine activity (Fig 2D, Table S5). Another low SI-upregulated module, F D1 M35, includes hub genes and enriched terms linked to dendrite organization and adhesion (Fig 2D, Table S5). In contrast, F D1 M31, downregulated in high SI, features genes enriched for cytoskeletal/membrane-related terms and hub genes in lipid trafficking, cytoskeletal organization, and growth factor signaling, suggesting transcriptional activity directed towards regulating structural processes (Fig 2D, Table S5). A similar male module, M D1 M22, is upregulated in low SI males, suggesting sex differences (Fig S5D, Table S6-7) [27]. These findings indicate increased neurite-forming activity in low SI females and reduced structural maintenance in high SI females.

Two D1- and three A2A-MSN modules include genes heavily involved in synaptic function. F D1 M40, downregulated low vs high SI and upregulated high SI vs controls, includes hub genes linked to synaptic vesicle sorting and release, with enrichment for protein processing- and transport-related terms (Fig 2D, Table S4-5). Similarly, F A2A M15, downregulated in low SI, includes genes involved in neurotransmitter/ neuropeptide vesicle release, with enrichment for terms involving protein localization and secretory granules (Fig 3D, Table S5). These results suggest reduced synaptic activity in low SI females across both subtypes, while increased D1-MSN synaptic activity is associated with high SI. The male module M D1 M10, also synaptic-related, is upregulated low SI vs control but downregulated in low SI vs high SI (Fig S5D, Table S6-7) [27]. F A2A M10, downregulated low SI vs controls/high SI, includes hub genes encoding receptors and components of receptor-mediated signal transduction. Enrichment results show significant association with terms involved in synaptic transmission and signal transduction (Fig 3D, Table S4-5). The male module M D2 M20 likewise includes genes involved in synaptic transduction however is upregulated in both low SI and high SI compared to control (Fig S7D) [27]. Overall, increased expression of genes involved in synaptic activity in D1-MSN correlates with high sociability in both sexes, with sex differences in genes involved in A2A-MSN synaptic input.

**Fig 3.**
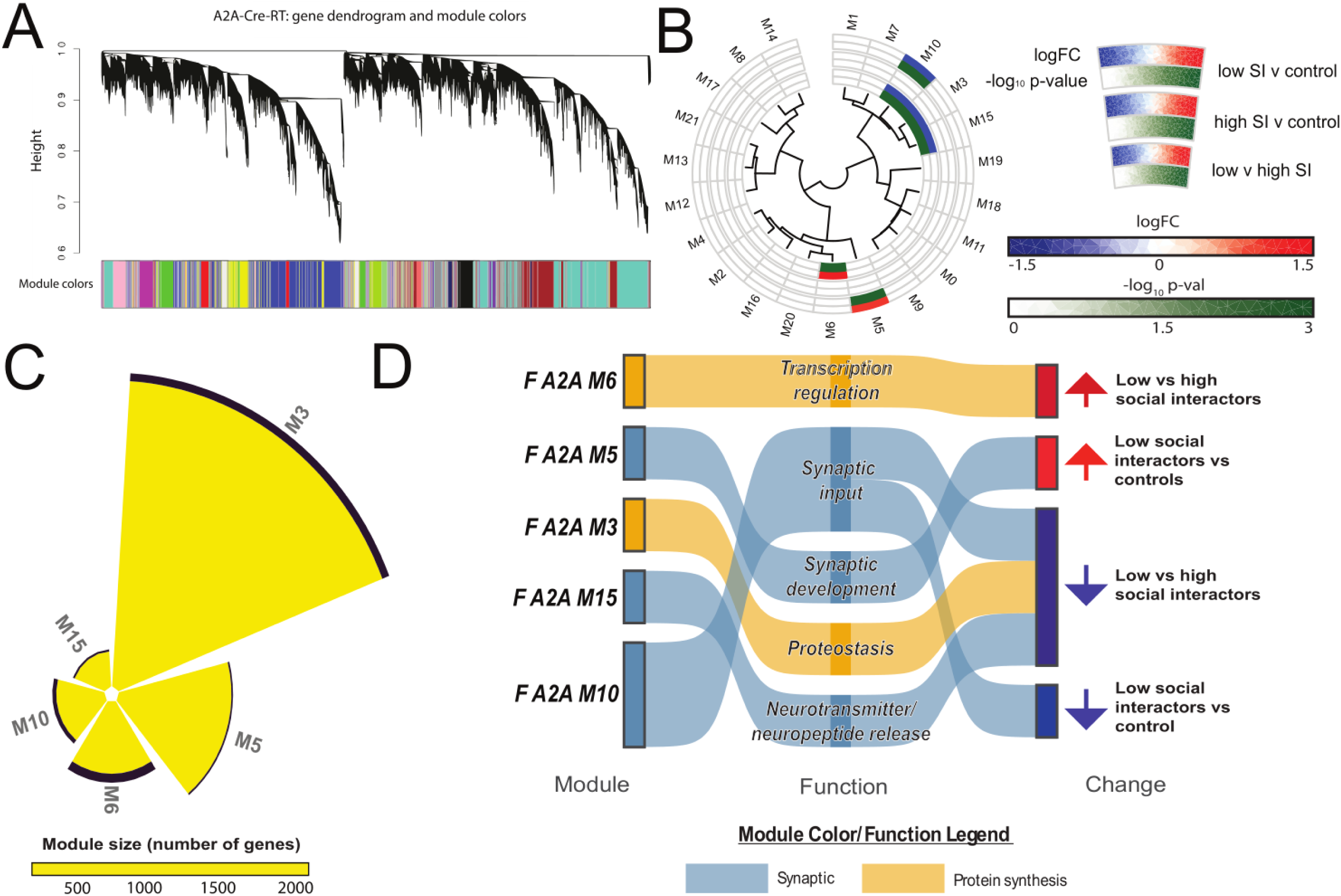
A) Dendrogram of genes from the female A2A-Cre-RT data with WGCNA module assignment represented in the colored bar below. B) Circos plots of the significant A2A-MSN modules with each ‘slice’ corresponding with one module. The plot is organized based on the module eigengene dendrogram, shown in the center, which positions related modules close to one another. Modules that aren’t significant are uncolored. C) Circular bar plots of A2A-MSN modules significantly associated with stress susceptibility and/or resilience in female socially stressed mice. The proportion of the module that includes differentially expressed genes from the A2A high SI vs low SI comparison is highlighted in purple. D) Sankey plots of significantly differentially regulated modules in the A2A-MSN of female socially stressed mice; modules are renamed based on their original WGCNA module number and functional categorization based on hub gene and enrichment analysis characterization.

Other significant synaptic modules include genes involved in synaptic maintenance. F D1 M41, linked to synaptic structural integrity and transport, is upregulated low vs high SI/controls (Fig 2D, Table S5). F A2A M5 includes hub genes involved in vesicle trafficking, encoding receptors/channels as well as synaptic structure formation (Fig 3D, Table S4-5). This module is also upregulated low SI vs controls, suggesting that genes involved in distinct aspects of synaptic maintenance are upregulated in low SI female mice by MSN subtype. In contrast, male modules that include genes involved in synaptic maintenance, M D1 M34 and M D1 M36, are downregulated low SI vs control (Fig S5D, Table S6-7) [27]. These results suggest that the regulation/modulation of synaptic activity may underlie sex differences.

F D1 M18 module, upregulated high SI vs control and downregulated low SI vs high SI, includes genes encoding mitochondrial respiratory chain and ribosomal (cytoplasmic and mitochondrial) subunits (Fig 2D, Table S4-5). F A2A M3 contains transcriptional/mitochondrial genes (Fig 3D, Table S4). These results indicate reduced mitochondrial activity in low SI D1-MSNs, with possible sex differences across MSN subtypes.

In human MDD WGCNA results, modules containing genes involved in synaptic transmission and regulation are upregulated in female controls, while a module with genes involved in synaptic development is downregulated relative to males. In MDD, synaptic development is downregulated in females. Mitochondrial and immune/inflammatory modules are upregulated in females in both control and MDD groups (Fig S9, Table S8-9).

### Female D1- and A2A-MSN modules with highest DEG proportions share common regulatory components

Modules with highest DEG proportions were selected for analysis. F D1 M12, upregulated low vs high SI, includes adhesion and cell-matrix interaction hub genes (Fig 4A, Table S4-5). DEGs include Vtn, KATP channel genes, and Lars2, all upregulated in low vs high SI. F A2A M10, downregulated low SI, includes downregulated DEGs involved in synaptic signaling (Fig 4B, Table S4-5). Enrichment analysis links both modules to PI3K-Akt-mTOR signaling (Fig 4C). Genes with high membership scores were analyzed for transcriptional regulation (Fig 4D). Nf1 and Klf6 emerged as key regulators. Nf1-regulated D1 M12 genes are linked to migration and growth factor signaling, A2A M10 genes to synaptic functions (Fig 4E).

**Fig 4.**
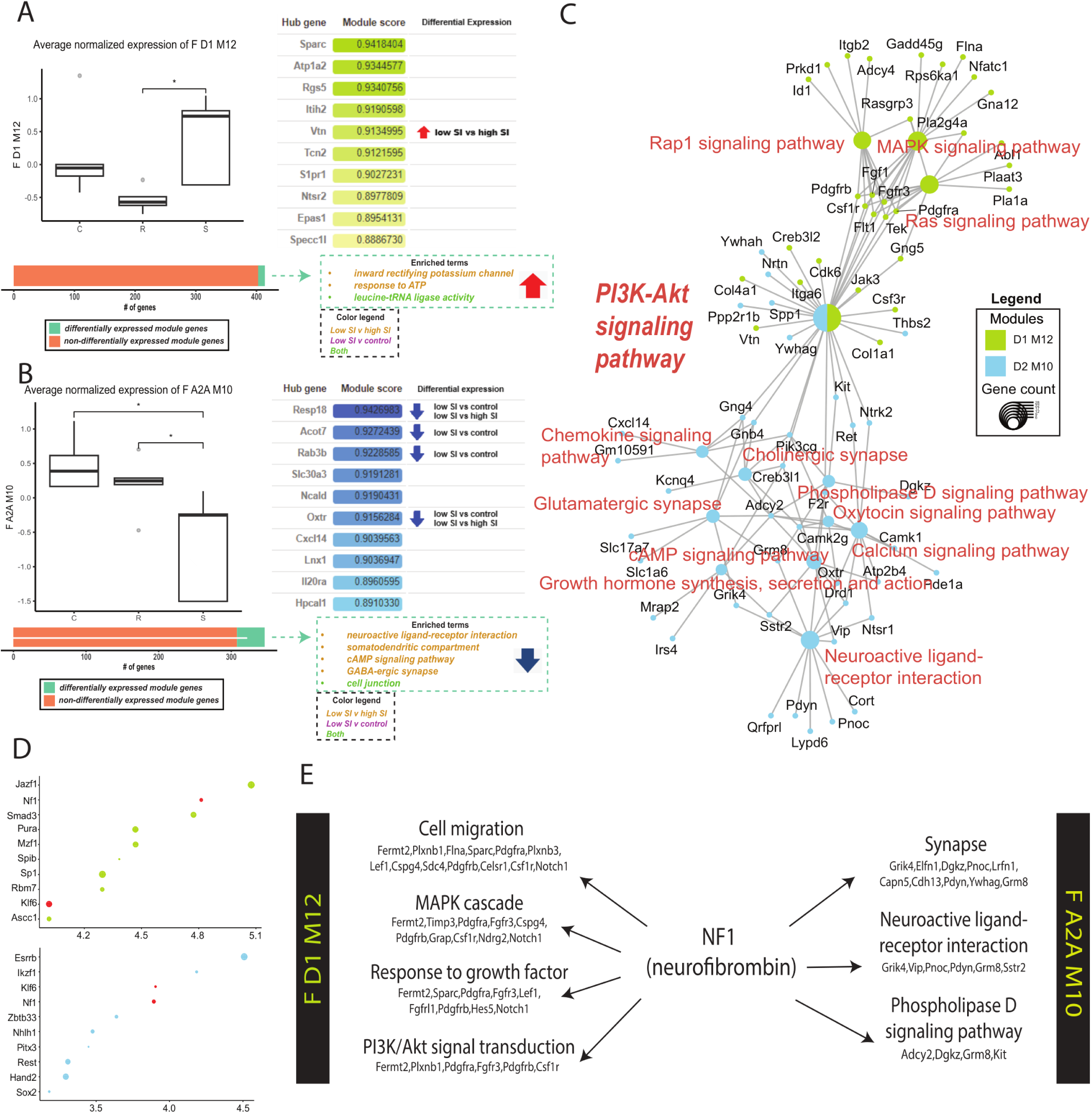
Subtype-specific modules with the largest proportion of differentially regulated genes between stress group selected for further analysis. A) (left, above) boxplot visualizing expression level of F D1 M12 by stress group (left, below) bar chart visualizing the proportion of genes in the D1 M12 module that are differentially regulated between the stress groups (right, above) top hub genes from D1 M12 with change indicated to the right (right, below) enrichment results for genes differentially expressed within the module. B) (left, above) box plot visualizing expression level of F A2A M10 by stress group (left, below) bar chart visualizing the proportion of genes in the D2 M10 module that are differentially regulated between the stress groups (right, above) top hub genes from A2A M10 with change indicated to the right (right, below) enrichment results for differentially expressed module genes. C) Gene-concept network plot visualizing D1 M12 and D2 M10 genes involved in the PI3K-Akt signaling pathway. D) (top) top transcription regulators identified using top genes from D1 M12 and (bottom) A2A M10. E) Enriched terms associated with NF1-regulated modular genes.

### Cross species consensus modules

Consensus module analysis was performed using the female CWDS dataset, male CSDS and human MDD datasets [23, 27]. 4029 genes overlapped across processed datasets (Fig 5A,B, Table S10). After module construction, relationship to experimental groups was assessed. One module, module 3, is differentially regulated across mice and humans (Fig 5C-E). Boxplots using the consensus module eigengenes indicate that module score medians all center around 0 for across all datasets with similar interquartile ranges (Fig S10). Hub genes involve transcription regulation and PI3K/Akt signaling (Table S10). Enrichment analysis shows that the module is involved in transport and synaptic development as well as inositol phosphate activity (Fig 5E, Table S11). Positive module score genes are enriched for synaptic activity and phosphatidylinositol-mediated signaling; negative module score genes for structural and translation terms.

**Fig 5.**
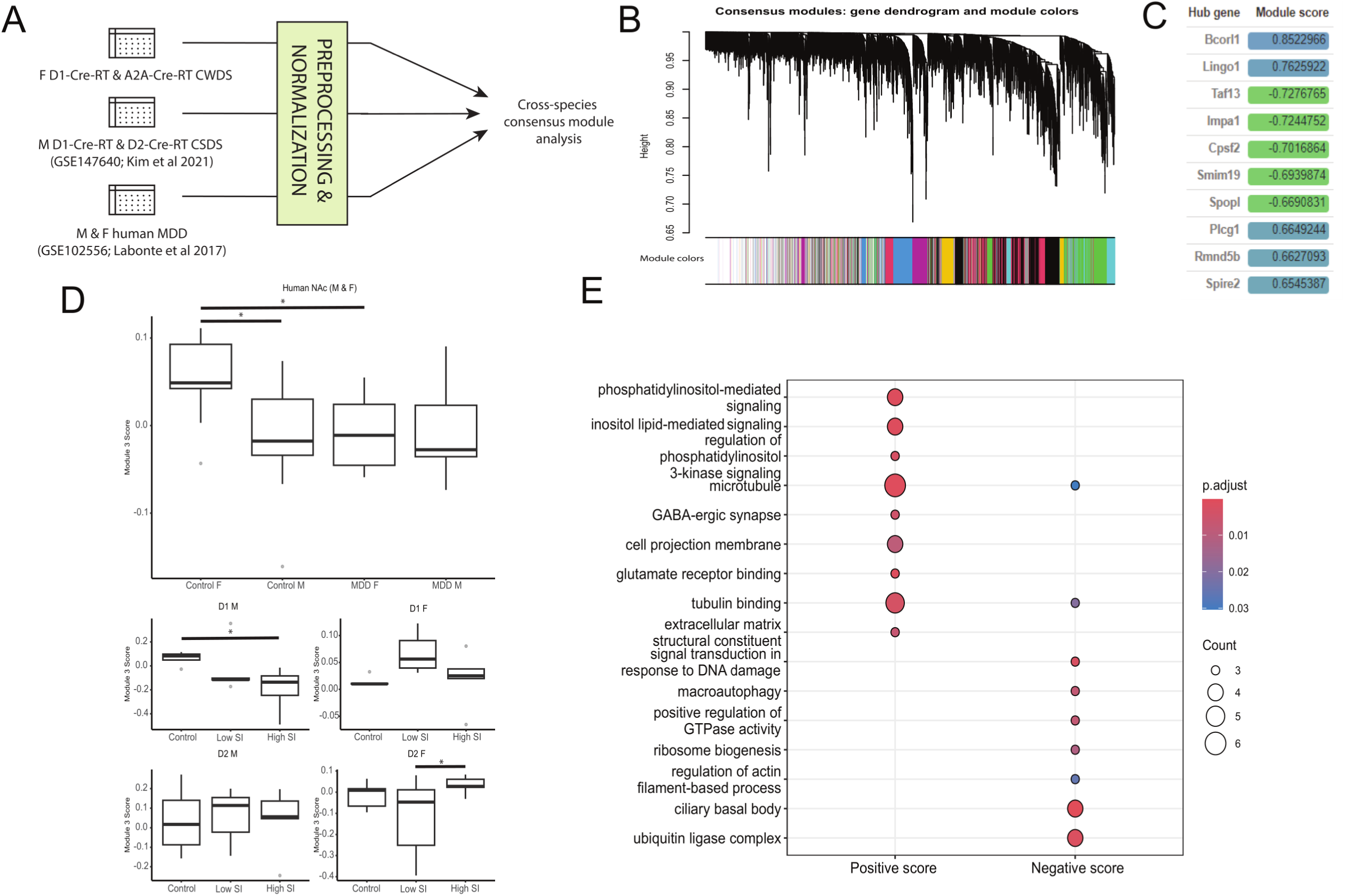
A) cross-species consensus module analysis was performed using transcriptomic data from male and female MDD NAc, D1- and D2-Cre-RT male mice that had undergone the chronic social defeat stress paradigm (CSDS) and the female CWDS dataset. B) Dendrogram of consensus module analysis with module assignment represented by the colored bars below. C) Top hub genes from consensus module three Network plot visualizing the consesus module 3. D) Boxplot of module 3 expression in (top) NAc of male and female human, (middle) D1-MSN of stressed male and female mice and (bottom) A2A-MSN of stressed male and female mice E) Enrichment analysis results of positive and negative consensus module 3-scored genes.

## Discussion

We investigated sex-specific effects of chronic social stress on NAc MSN subtype transcriptomes. Using female D1-Cre-RT and A2A-Cre-RT mice exposed to CWDS, we identified unique DEGs and co-expression networks linked to social interaction phenotype, as well as networks showing sex-specific regulation in mice and humans. Genes upregulated in D1-MSNs of low-SI females suggest altered respiratory activity. Two encode ATP-sensitive potassium channels (KATP), which regulate excitability based on ATP levels, and *Lars2* encodes a mitochondrial aminoacyl-tRNA synthetase. KATP expression increases under hypoxia, while altered mt-aaRS expression is linked to mitochondrial dysfunction [28, 29]. These findings align with mitochondrial changes in D1-MSNs associated with depressive-like behavior and social deficits [30, 31]. Together, the DEGs and module results suggest mitochondrial dysfunction contributes to low SI in females, while mitochondrial genes and biogenesis are increased in high SI females. WGCNA also shows downregulation of co-expression networks for energy-demanding processes like protein synthesis, transport, and synaptic release in low-SI D1-MSNs. A recent study found that chronic dietary supplementation of urolithin A, a gut metabolite known to enhance mitochondrial health, produces anxiolytic effects in high anxiety mice mediated by the reversal of dysfunctional mitochondrial mechanisms in the MSNs of NAc [40]. Given the high degree of comorbidity between anxiety disorders and mood disorders, these results suggest that targeting mitochondria in MSN subtypes may produce therapeutic effects.

There were far more DEGs in female A2A-MSNs relative to D1-MSNs. In low-SI females, top upregulated A2A-MSN genes are involved in structure and transcription, while top downregulated genes relate to synaptic activity, consistent with enrichment analysis results. Our previous studies observed male low-SI mice display increased frequency of excitatory input and number of thin and stubby spines on A2A-MSNs [30, 32]. This suggest sex differences A2A-MSN synaptic activity, with reduced activity in low-SI females potentially driving stress susceptibility. Supporting this, WGCNA identified multiple A2A-MSN gene co-expression networks impacted by chronic social stress that are composed of genes involved in synaptic processes. Although the modules with genes linked to synaptic input and release are downregulated in low-SI female mice, modules linked to synapse development/formation is upregulated, possibly as a compensatory or adaptive mechanism.

We identified sex differences by comparing female CWDS results with male CSDS modules [27]. In D1-MSNs, differences were observed in structural regulation modules: F D1 M31 is downregulated in high SI females, whereas M D1 M22 is upregulated in low SI males. Both modules include genes involved in cytoskeletal regulation and lipid processing; however, F D1 M31 genes involve growth factor signaling, whereas M D1 M22 genes involve lipid processing. These results suggest stress may impair growth factor– dependent cytoskeletal remodeling in females, limiting dendritic expansion and plasticity, while in males, altered lipid metabolism may affect membrane composition and spine stability. In A2A-MSNs, there are differences in genes involved in synaptic input. F A2A M10 is downregulated in low SI vs control/high-SI females while M D2 M20 is upregulated in low-SI males. Although only a few genes overlap, hub genes in both regulate GPCR signaling, vesicle transport, and cytoskeletal dynamics. Sex differences in MSN morphology and connectivity have been reported; though the mechanism remains unclear [33]. These results suggest stress susceptibility may be the result of opposing transcriptional trajectories: reduced GPCR activity produces dampened response in low-SI female A2A-MSNs, while male low-SI A2A-MSNs may experience enhanced maladaptive excitatory signaling. Further research is needed to clarify A2A-MSN input alterations.

In D1-MSNs, the D1 M12 module contains the highest proportion of DEGs, with upregulation in low vs high SI groups. Multiple hub genes encode proteins involved in migration. Vitronectin, a glycoprotein regulating cell adhesion and migration, is upregulated in low SI females [34]. Other hub genes are involved in GPCR signaling and homeostasis. Stress-induced depressive behaviors are linked to D1-MSN dendritic atrophy via GTPase RhoA upregulation [32]. Given its involvement in GPCR signaling and cytoskeletal regulation, D1 M12 likely contributes to this atrophy in low-SI mice.

Among A2A-MSN modules, A2A M10 had the highest proportion of DEGs. Its hub genes, involved in synaptic and GPCR signaling, include *Oxtr* and *Resp18*, both downregulated in low-SI mice, suggesting reduced peptidergic signaling [35]. *Oxtr* encodes the oxytocin receptor, linked to social reward and expressed in both D1- and A2A-MSNs [36]. All low vs. high SI DEGs in A2A M10 were downregulated and enriched for synaptic terms. The module itself was downregulated in low-SI vs. controls and high-SI females. Prior studies show increased excitatory input onto A2A-MSNs in low-SI mice post-defeat [20], suggesting reduced A2A M10 activity reflects weakened modulatory synaptic input.

Enrichment analysis revealed that both D1 M12 and A2A M10, oppositely regulated in low SI, are significantly enriched for the PI3K-Akt pathway, involved in cell growth and survival [37]. It also plays a neuroprotective role under cellular stress [38]. The PI3K-Akt genes in D1 M12 are associated with the MAPK/ERK pathway. Dendrite morphogenesis involves both the PI3K and the MAPK pathway, and PI3K-dependent dendritic remodeling plays a neuroprotective role [39, 41]. The D2 M10 genes linked to PI3K/Akt center on synaptic signal transduction. This is consistent with previous studies which have established that the PI3K/Akt pathway is involved in various components of synaptic activity [37].

Consensus module analysis of the female CWDS, male CSDS, and human MDD datasets identified module 3 as sex-specific in both mice and humans [23, 27]. Results show an upregulation in module 3 activity in the female human NAc at baseline that is significantly reduced in the MDD condition. Additionally, the cell subtype-specific mouse transcriptomes display a reduction in module 3 activity in the A2A-MSN of low SI females. This suggests that module 3 in A2A-MSNs represents a sex-specific mechanism that may be reduced in NAc A2A-MSNs of human females with MDD. Conversely, high SI males show reduced module 3 activity in D1-MSNs, suggesting that the reduction of module 3 activity in D1-MSNs of male mice may be protective. Further work is required to determine whether baseline sex differences in MSN subtypes drive sex- and subtype-specific PI3K-Akt responses. Hub genes in this module regulate transcription and PI3K-Akt signaling, while enrichment analysis indicates involvement in mTOR signaling, downstream of PI3K-Akt. Abnormal PI3K-Akt pathway activity in the NAc is linked to multiple substance use disorders [42–44]. Some studies suggest sex-specific involvement in the NAc [45–46]. A recent study investigating ketamine’s anti-depressant effects observed sex-specific AKT-mTOR signaling alterations in the hippocampus; decreased activity in males linked to memory impairment, increased activity in females linked to resilience [47]. Although further research is required, our findings suggest PI3K/Akt signaling is involved in the sex-specific stress response in the NAc in a subtype-specific manner.

iRegulon analysis identified key transcriptional regulators, notably NF1 (Neurofibromin 1), as a shared regulator of F D1 M12 and F A2A M10 modules (Fig 4D). NF1, a tumor suppressor, influences migration, cytoskeletal dynamics, and stress responses through pathways including Ras/MAPK, Raf/MEK/ERK, PI3K/AKT/mTOR, and cAMP/PKA (Fig 4E) [48]. It also contributes to reward and motor behavior via its role in D1- and A2A-MSNs and is linked to striatal neuropsychiatric conditions [49–50]. NF1 may act as a signaling hub for GPCR pathways and could mediate stress-induced effects on MSN subtypes via PI3K/AKT/mTOR signaling.

There were a few limitations to the study. To begin with, there was a small number of DEGs identified in the D1-MSN of female mice. This may have been the result of using RiboTag-based approach which may yield more conservative DEG profiles compared to total RNA-seq. Additionally, the pooling of animals may have limited the estimation of inter-individual variance, which could have reduced the power to detect subtle differential expression. Moreover, estrous cycle stage was not assessed, limiting our ability to determine how fluctuations in ovarian hormones such as estrogen may influence nucleus accumbens translatomic profiles in female mice [51–52]. Future studies incorporating estrous cycle staging will be important for disentangling hormone-dependent transcriptional mechanisms underlying stress phenotype in females. In addition to this, due to the lack of available transcriptomic data from male mice performing CWDS, our consensus module analysis relied on male CSDS data. Although the difference in stress type may influence outcomes, both paradigms involve social stress but differ in physical versus vicarious exposure. Given that the purpose of WGCNA is to find conserved networks and test how they differ by stress and sex, our results allow us to examine both conserved and paradigm-specific transcriptional modules. Future studies should examine the transcriptomic alterations induced in MSN subtypes by CSDS, including newly developed female models, relative to CWDS, to investigate how different forms of stress contribute to MDD onset. In addition to this, further work will be needed to experimentally validate stress-associated genes and network-level signatures identified in this study using targeted molecular approaches. An important aspect of our study is the analysis of female subjects, extending beyond prior transcriptomic studies that focused solely on males [53–55].

In conclusion, chronic stress affects MSN subtypes in a cell-type- and sex-specific manner. Morphological, synaptic, mitochondrial, and transcriptional changes contribute to female vulnerability, potentially driven by PI3K/Akt/mTOR signaling and NF1. By identifying sex-specific molecular signatures in NAc MSN subtypes after social stress, this study offers key insights into MDD mechanisms and highlights potential targets for sex-informed therapies.

## Supporting information

Supplemental Figures (all)

Supplemental Table titles & descriptions

## Author contribution

Conceptualization, MEF, MKL, GK; Methodology, MEF, MKL, SA, GK; Formal analysis, GK, MB; Investigation, MEF, DF, JO; Resources, MKL; Writing – Original Draft, GK, RC, MKL; Reviewing & Editing, MKL, RC, MEF, SAA; Visualization, GK; Funding Acquisition, MEF, MKL

