## Supplemental Figures (all) for "Nucleus accumbens neuron subtype translatome signatures in socially stressed females"

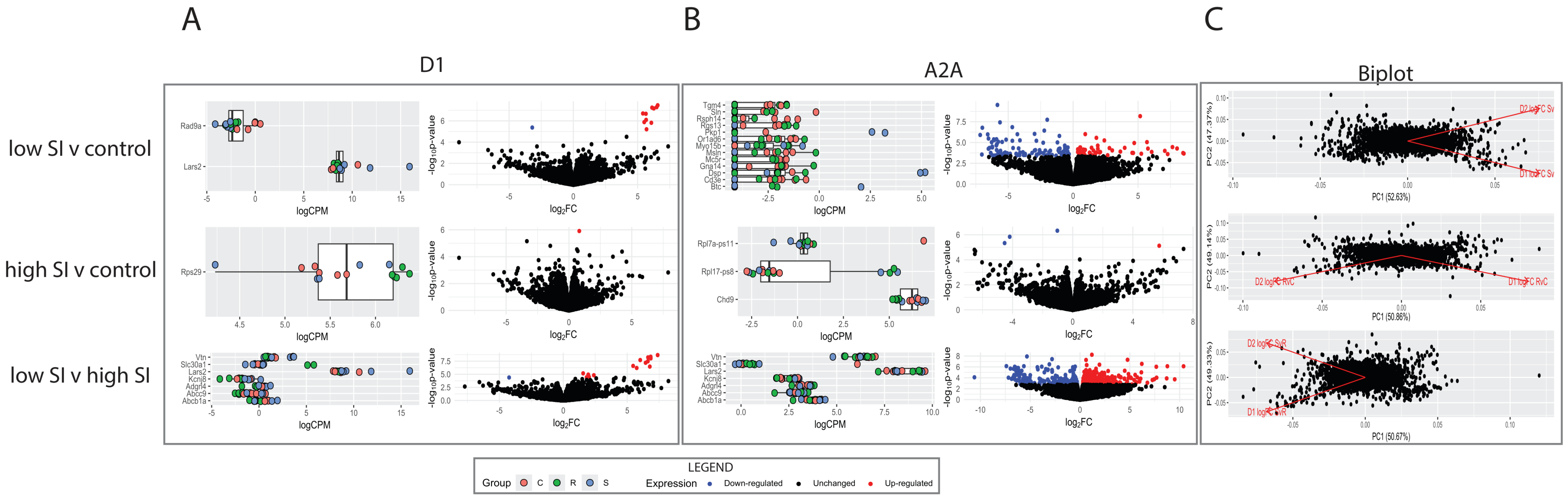


Supp Fig 1. Boxplots visualizing normalized expression of top DEGs (as indicated in Figure 1) by group and volcano plots visualizing expression change and statistical significance for each comparison indicated on the far left side with DEGs highlighted in A) D1-MSNs and B) A2A-MSNs. C) Biplots visualizing relationship between the effect of stress phenotype membership in D1-MSNs and D2-MSNs.

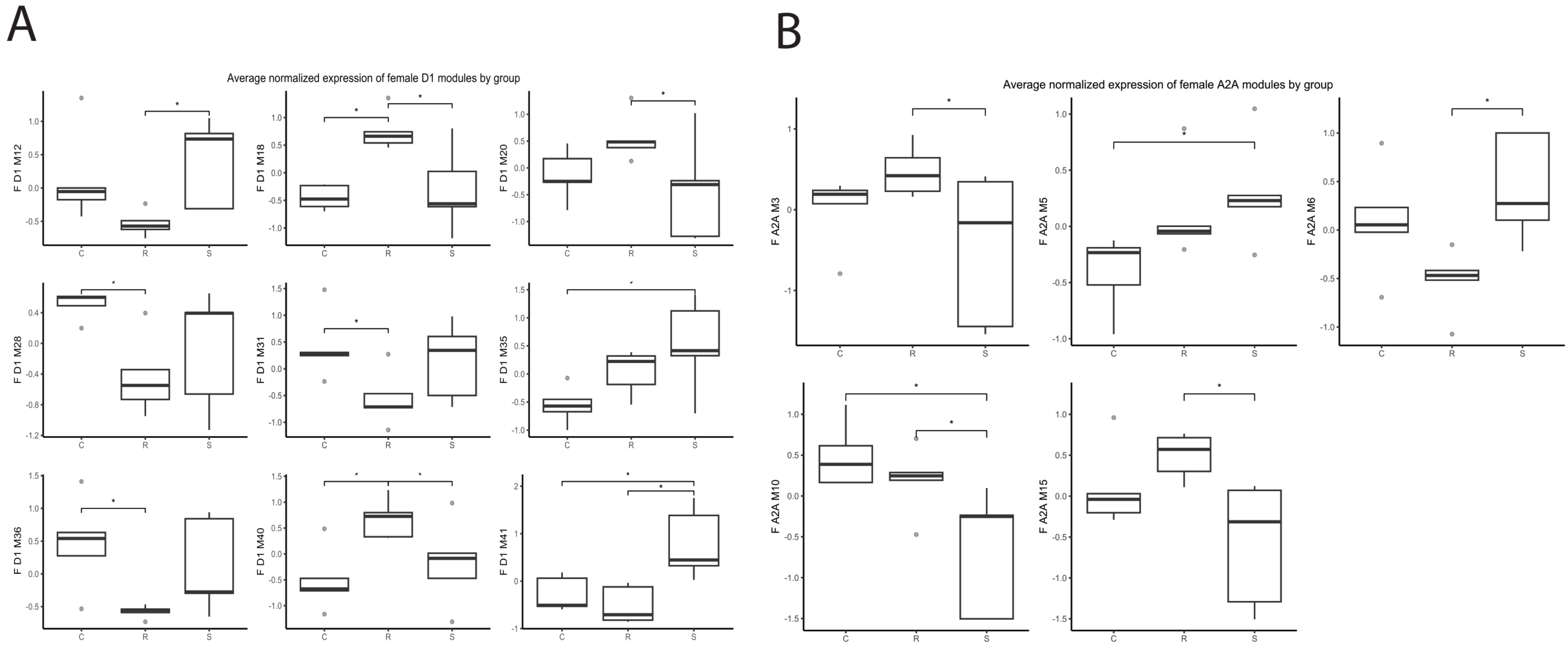


Supp Fig 2. boxplots visualizing the average normalized expression level by stress phenotype of significant gene co-expression modules identified in A) D1-MSNs of female CWDS mice, B) D2-MSNs of female CWDS mice

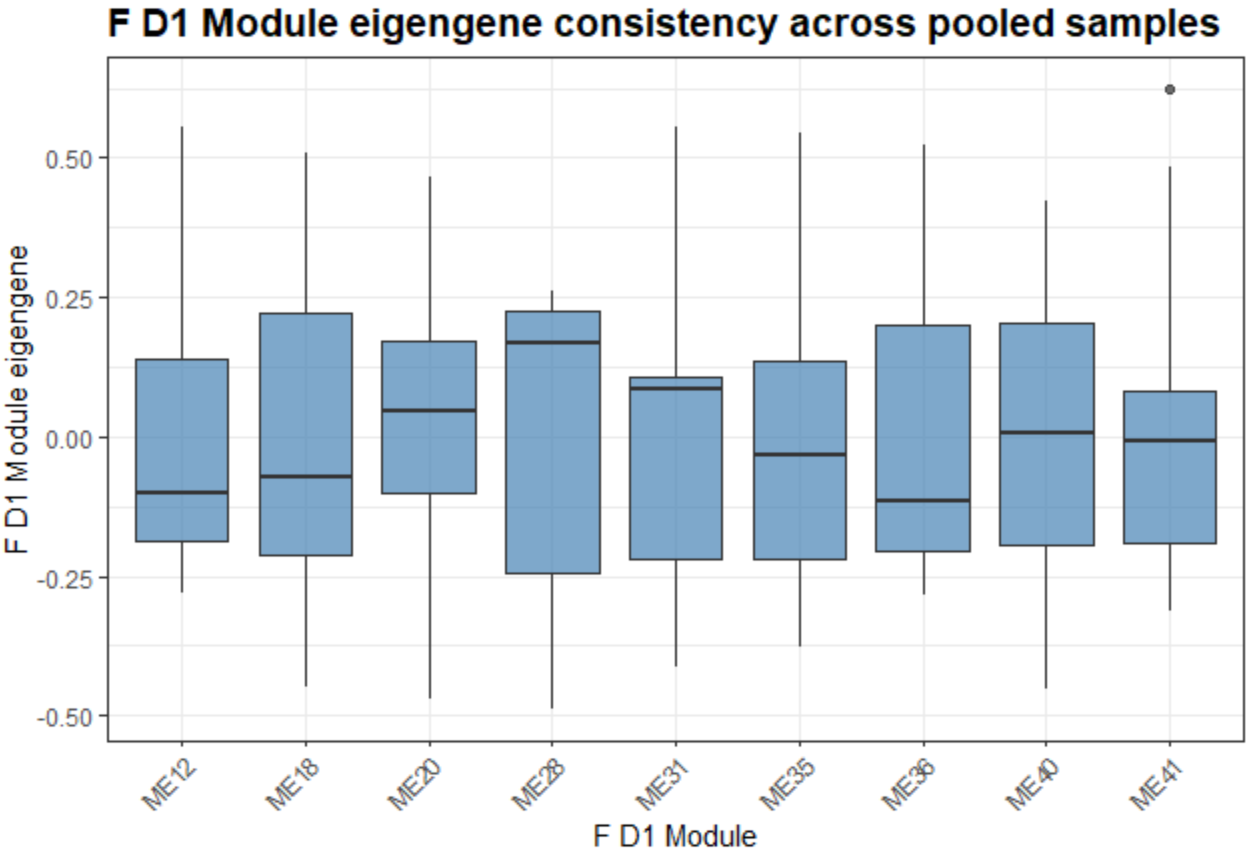


Supp Fig 3. Boxplots visualizing module eigengene expression levels across female D1-Cre-RT pooled samples

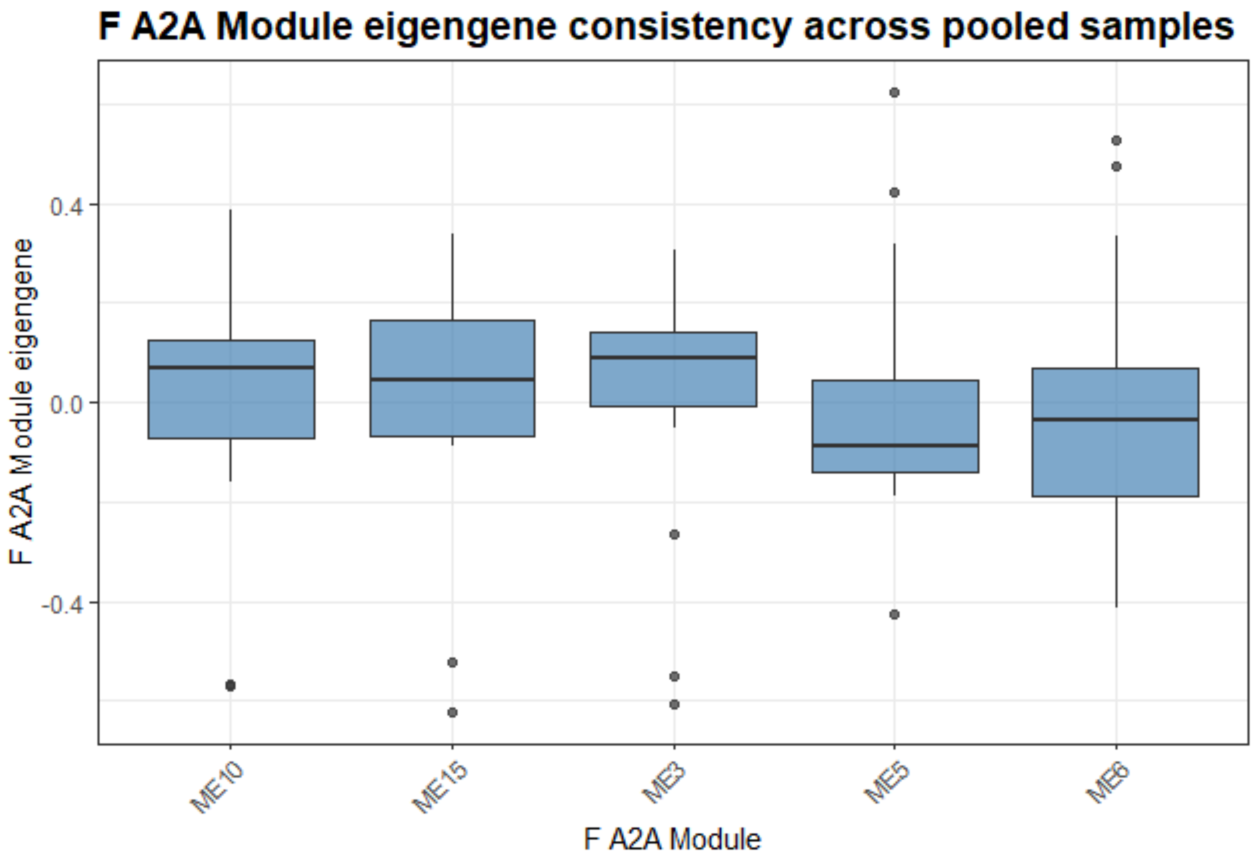


Supp Fig 4. Boxplots visualizing module eigengene expression levels across female A2A-Cre-RT pooled samples

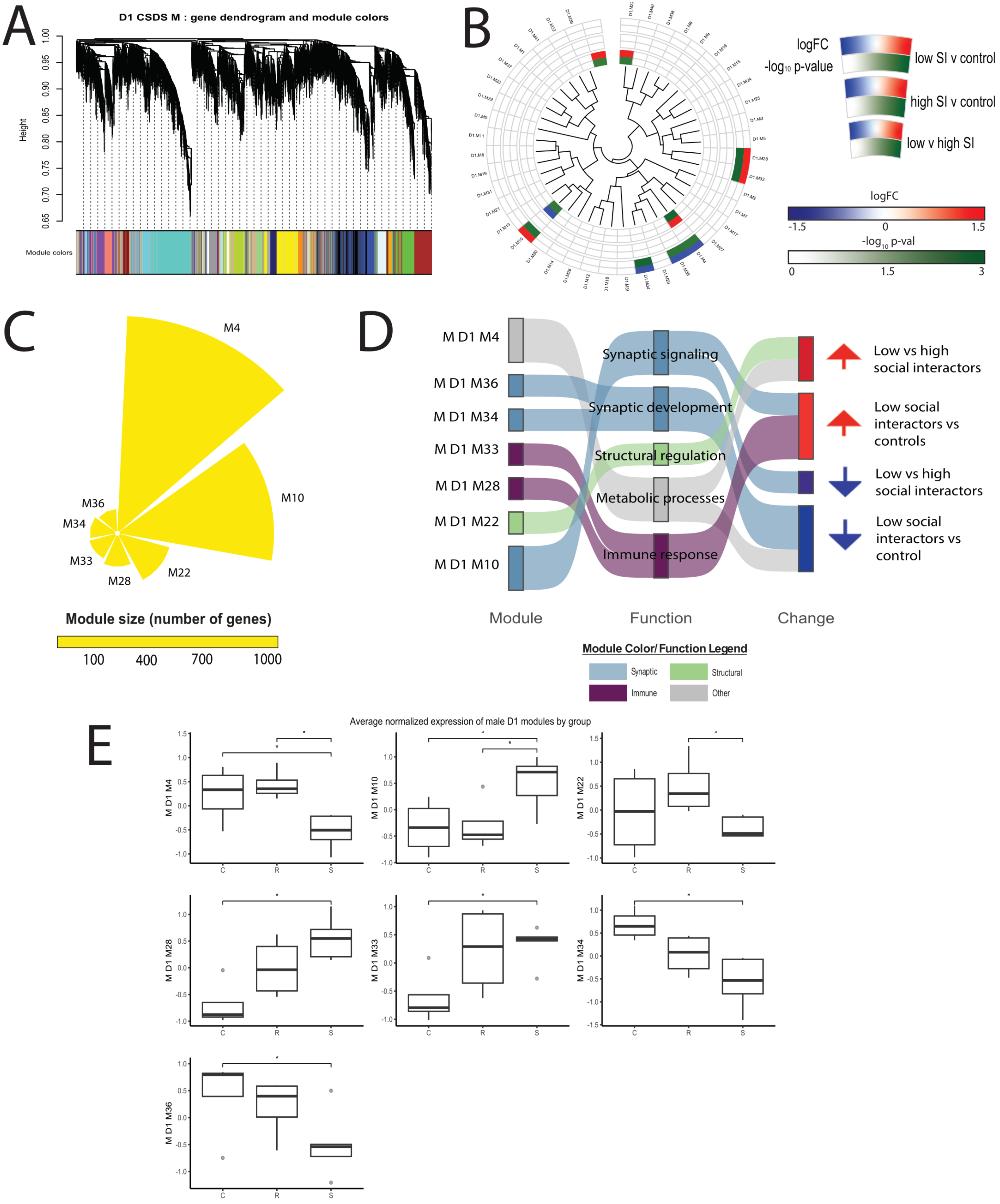


Supp Fig 5. A) Dendrogram of genes from the male D1-Cre-RT data with WGCNA module assignment represented in the colored bar below. B) Circos plots of the significant male D1-MSN modules with each 'slice' corresponding with one module. As indicated in the legend to the right, the outermost three wedges represent the log fold change in module expression levels for each stress group comparison while the innermost three slices represent the –log10 p-value for the respective comparisons. Modules that aren't significant are uncolored. C) Circular bar plots of male D1-MSN modules significantly associated with stress susceptibility and/or resilience in male socially stressed mice. D) Sankey plots of significantly differentially regulated modules in the D1-MSN of male socially stressed mice; modules are renamed based on their original WGCNA module number and functional categorization based on hub gene and enrichment analysis characterization.

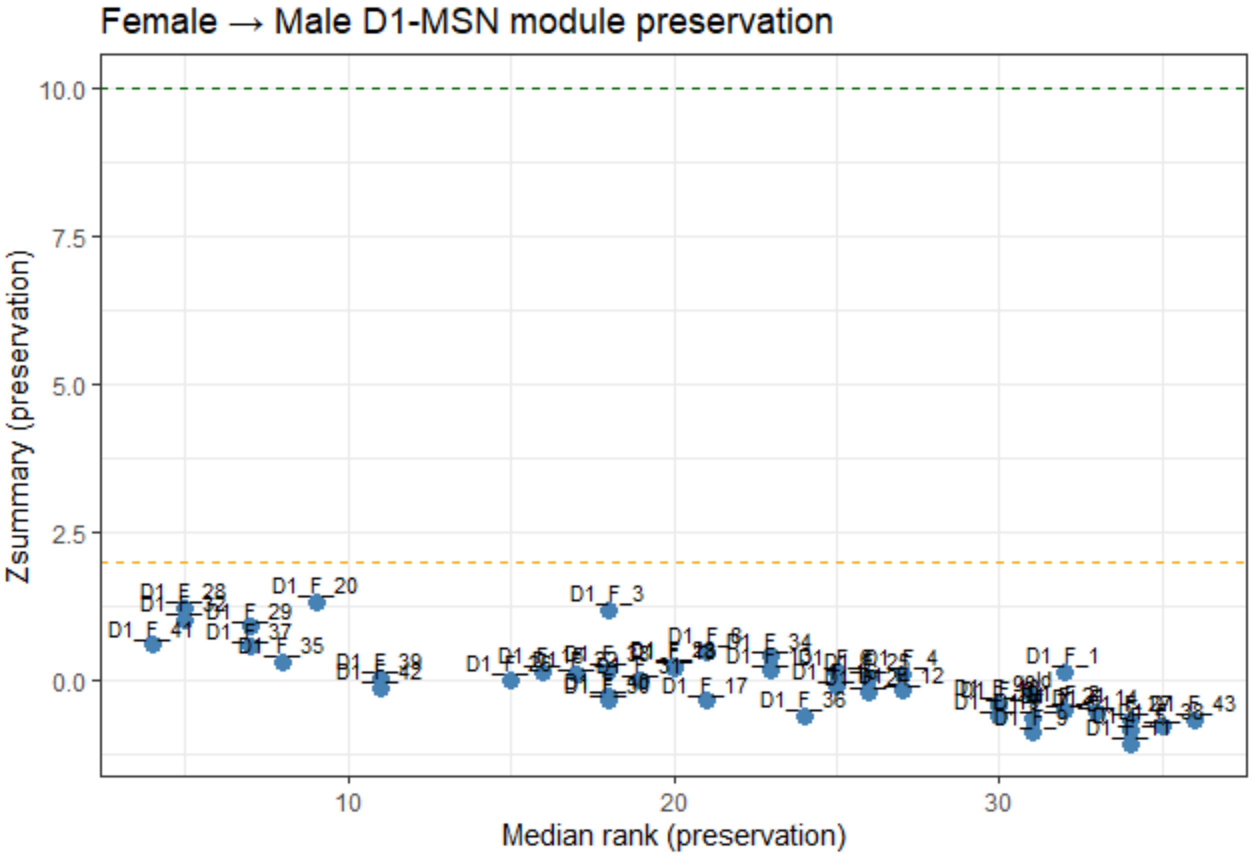


Supp Fig 6) Module preservation analysis of female to male D1-MSNs. Each point represents a co-expression module defined in female D1-MSNs. Zsummary values on the y-axis indicate the degree of preservation in male D1-MSNs, with higher values reflecting stronger preservation.

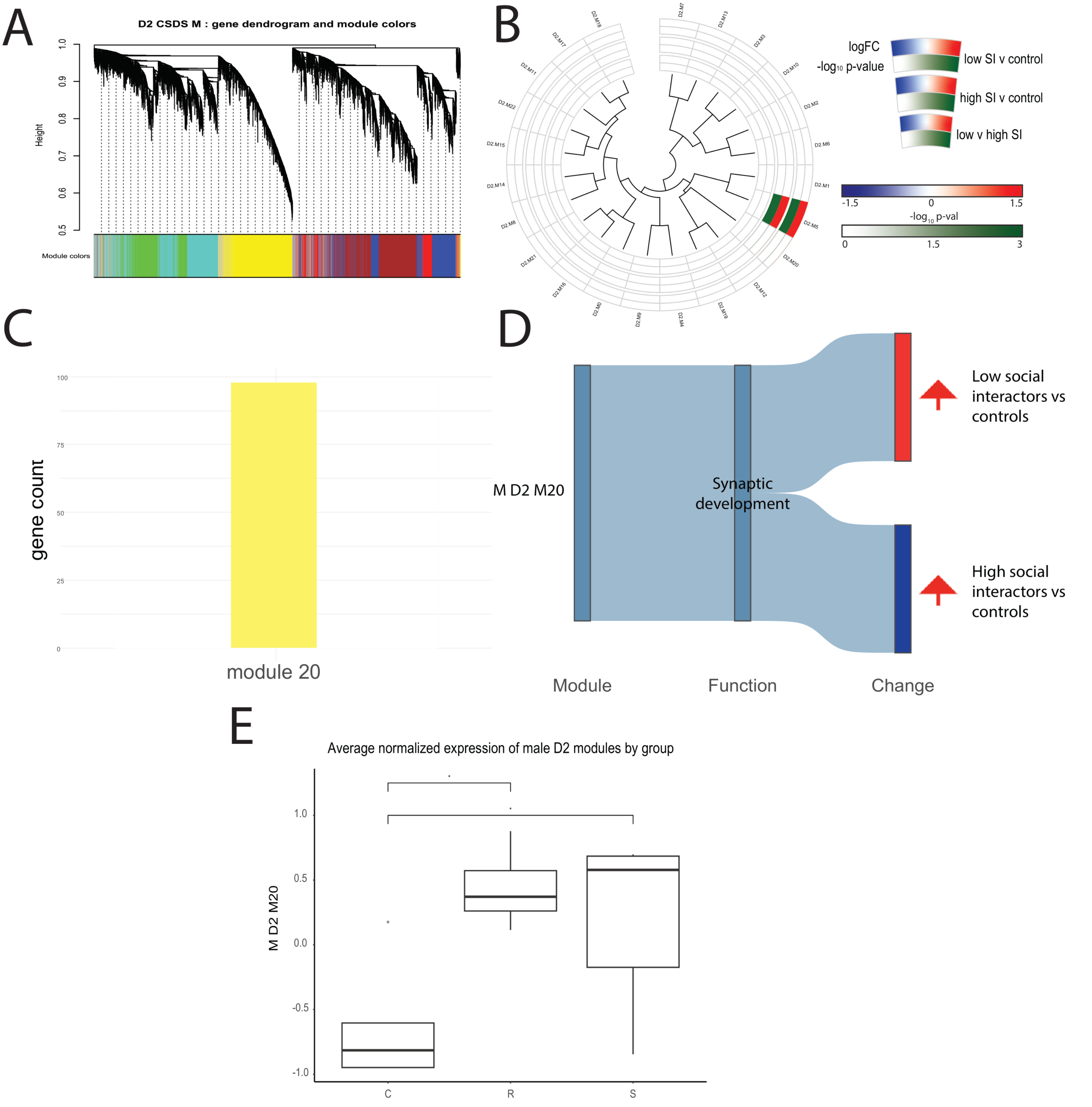


Supp Fig 7. A) Dendrogram of genes from the male D2-Cre-RT data with WGCNA module assignment represented in the colored bar below. B) Circos plots of the significant male D1-MSN modules with each 'slice' corresponding with one module. As indicated in the legend to the right, the outermost three wedges represent the log fold change in module expression levels for each stress group comparison while the innermost three slices represent the –log10 p-value for the respective comparisons. Modules that aren't significant are uncolored. C) Bar plot of male D2-MSN module significantly associated with stress susceptibility and/or resilience in socially stressed mice. D) Sankey plots of significantly differentially regulated modules in the D1-MSN of male socially stressed mice; modules are renamed based on their original WGCNA module number and functional categorization based on hub gene and enrichment analysis characterization. E) Boxplots visualizing the average normalized expression level by stress phenotype of significant gene co-expression modules identified in D2-MSNs of male CSDS mice.

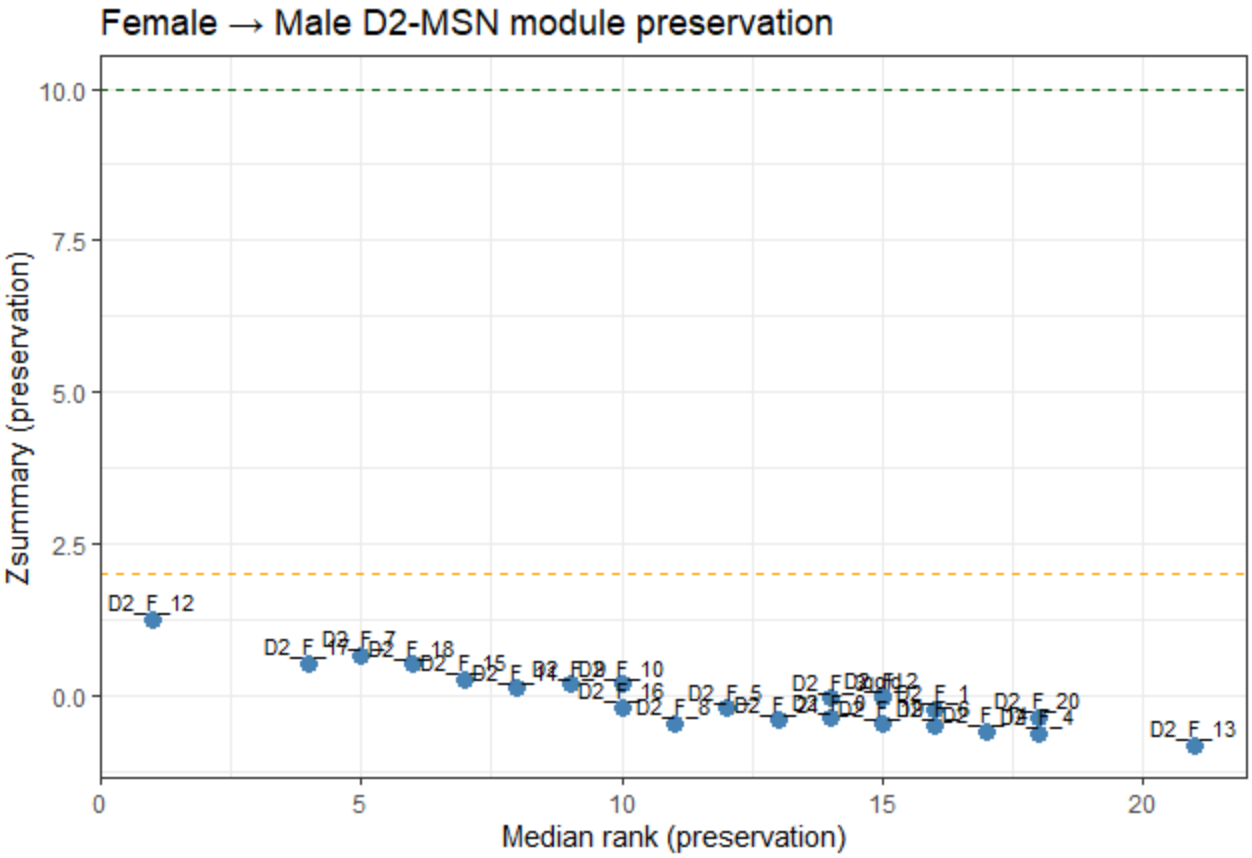


Supp Fig 8) Module preservation analysis of female to male D2-MSNs. Each point represents a co-expression module defined in female A2A-MSNs. Zsummary values on the y-axis indicate the degree of preservation in male D2-MSNs, with higher values reflecting stronger preservation.

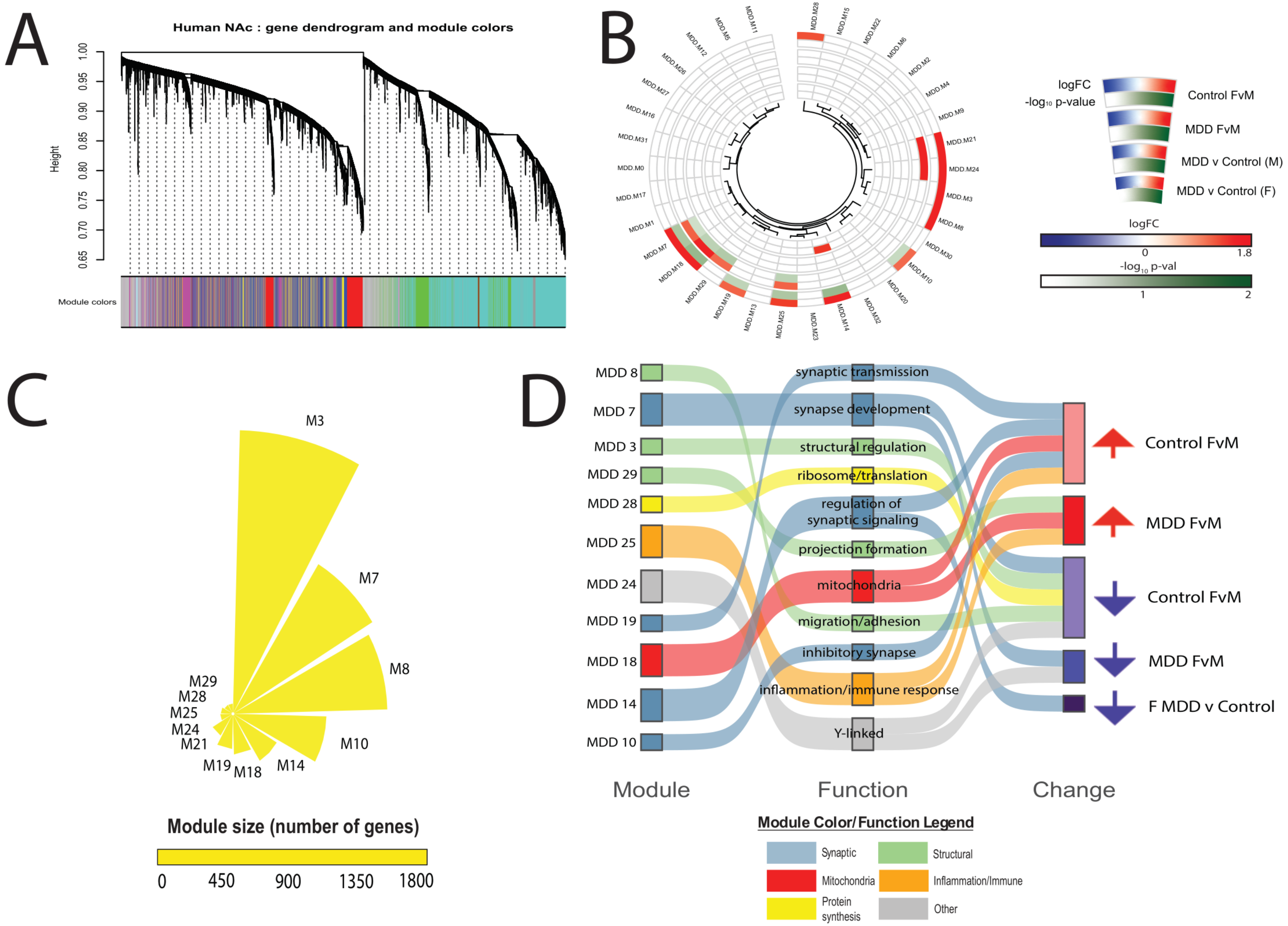


Supp Fig 9. A) Dendrogram of genes from the human MDD data with WGCNA module assignment represented in the colored bar below. B) Circos plots of the significant NAc modules with each 'slice' corresponding with one module. As indicated in the legend to the right, the outermost three wedges represent the log fold change in module expression levels for each stress group comparison while the innermost three slices represent the –log10 p-value for the respective comparisons. Modules that aren't significant are uncolored. C) Circular bar plots of human NAc modules significantly associated with stress susceptibility and/or resilience or with sex or both. D) Sankey plots of significantly differentially regulated modules in the human NAc; modules are renamed based on their original WGCNA module number and functional categorization based on hub gene and enrichment analysis characterization.

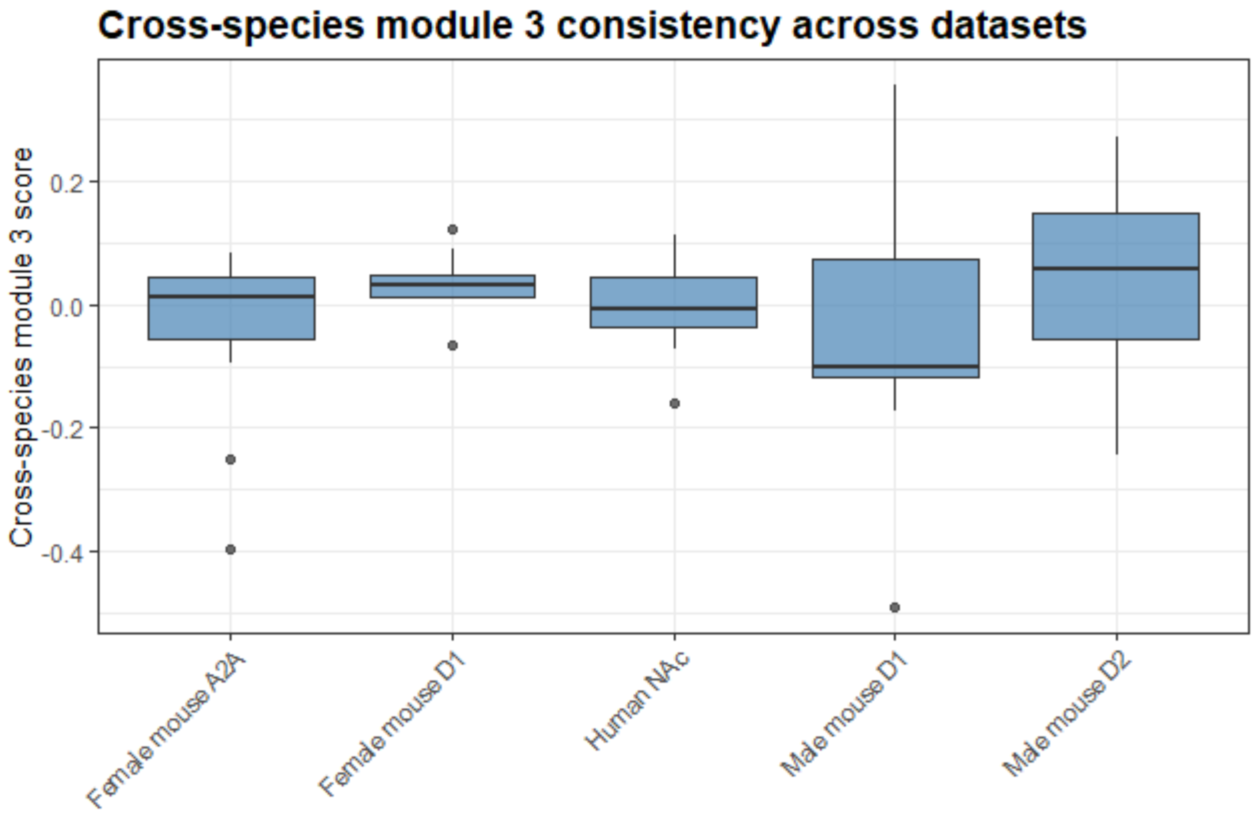


Supp fig 10. Boxplots visualizing cross-species module eigengene expression levels across male and female D1- and A2A-Cre-RT mice and human subjects
