## Supplemental Table titles & descriptions for "Nucleus accumbens neuron subtype translatome signatures in socially stressed females"

**Supplementary Table S1. Differentially expressed genes associated with social interaction phenotypes in female CWDS mice**

**Description:**
Differentially expressed genes (DEGs) identified from comparisons between social interaction (SI) groups in female mice subjected to the chronic witness defeat stress (CWDS) paradigm. Separate worksheets report results for D1-MSNs and D2-MSNs. Columns include Ensembl gene ID (ENSEMBL), gene symbol (SYMBOL), log2 fold change (logFC), nominal p-value (pvalue), adjusted p-value (padj), and the specific comparison performed (comparison; e.g., high SI vs low SI, high SI vs control).

**Supplementary Table S2. Functional enrichment analysis of DEGs using adjusted p-value thresholds**

**Description:**
Functional enrichment results for DEGs identified in female CWDS mice using adjusted p-value cutoffs. Separate worksheets correspond to individual cell type– and comparison–specific DEG sets and enrichment tools, including clusterProfiler and gprofiler2. clusterProfiler outputs include ontology, term ID and description, enrichment statistics, and contributing genes, while gprofiler2 outputs include functional term identifiers, source databases, term and query sizes, intersection statistics, and enrichment p-values.

**Supplementary Table S3. Functional enrichment analysis of DEGs using nominal p-value thresholds**

**Description:**
Functional enrichment results generated using gprofiler2 for DEG sets defined by a nominal p-value cutoff ($p < 0.05$). Results are reported in a single worksheet, with enrichment terms annotated by source database, statistical significance, term and query sizes, and gene intersections. The comparison column specifies the corresponding cell type, SI comparison, and direction of differential expression (e.g., D1 low SI vs control, upregulated).

**Supplementary Table S4. Functional enrichment of significant WGCNA modules in female CWDS datasets**

**Description:**
Functional enrichment analyses for WGCNA modules significantly associated with SI group in female CWDS datasets. Separate worksheets report enrichment results generated using clusterProfiler, gprofiler2, or DAVID for individual D1- and D2-MSN modules. Output columns reflect tool-specific formats, including ontology and functional category annotations, enrichment statistics, and contributing genes.

**Supplementary Table S5. Module membership and eigengene-based gene connectivity in female CWDS WGCNA**

**Description:**
Gene-level module membership information from WGCNA performed on female CWDS datasets. Columns include Ensembl gene ID, gene symbol, gene biotype, D1- and D2-MSN module assignments, and module membership (MM) values reflecting eigengene-based connectivity for each significant D1- and D2-MSN module.

**Supplementary Table S6. Functional enrichment of significant WGCNA modules in male CSDS datasets**

**Description:**
Functional enrichment results for significant WGCNA modules identified in male chronic social defeat stress (CSDS) datasets. Enrichment analyses were performed using clusterProfiler, gprofiler2, and DAVID, with separate worksheets corresponding to individual modules and tools. Reported columns include functional annotations, enrichment statistics, and gene-level contributions as defined by each method.

**Supplementary Table S7. Module membership and eigengene-based gene connectivity in male CSDS WGCNA**

**Description:**
Gene-level module membership information from WGCNA performed on male CSDS datasets. Columns include Ensembl gene ID, gene symbol, chromosomal location, gene biotype, gene description, D1- and D2-MSN module assignments, and module membership (MM) values for significant modules.

**Supplementary Table S8. Gene membership of significant human nucleus accumbens WGCNA modules**

**Description:**
Gene membership information for WGCNA modules significantly associated with major depressive disorder (MDD) in human nucleus accumbens (NAc) transcriptomic datasets. Columns indicate gene identifiers, module assignments, and module eigengene-based connectivity values for each significant module.

**Supplementary Table S9. Functional enrichment of significant human nucleus accumbens WGCNA modules**

**Description:**
Functional enrichment results for significant human NAc WGCNA modules. Enrichment analyses were conducted using clusterProfiler, gprofiler2, and DAVID, with tool-specific

output formats reported in separate worksheets. Results include functional annotations, enrichment statistics, and contributing genes.

**Supplementary Table S10. Gene membership of cross-species consensus WGCNA modules**

**Description:**
Gene membership information for cross-species consensus WGCNA modules derived from integrated mouse and human transcriptomic analyses. Columns indicate gene identifiers, module assignments, and module-specific connectivity measures used in consensus network construction.

**Supplementary Table S11. Functional enrichment of cross-species consensus WGCNA modules**

**Description:**
Functional enrichment analyses of cross-species consensus WGCNA modules. Enrichment results were generated using established functional annotation tools and are reported in tool-specific formats across worksheets, including functional categories, statistical significance measures, and contributing gene sets.
